# Remote focusing system for simultaneous dual-plane mesoscopic multiphoton imaging

**DOI:** 10.1101/503052

**Authors:** Dmitri Tsyboulski, Natalia Orlova, Fiona Griffin, Sam Seid, Jerome Lecoq, Peter Saggau

## Abstract

We present a dual-plane mesoscopic imaging system capable of simultaneous image acquisition from two independent focal planes. The system was designed as an add-on to a recently introduced large field-of-view two-photon microscopy system, developed by Sofroniew, *et al*., eLife, 5, e14472, 2016. In this work, we merge two advanced multiphoton imaging technologies, *i.e*., temporal-division multiplexing and remote focusing, to maintain diffraction-limited resolution at both imaging planes, and achieve a more than 2-fold increase in the system’s overall imaging throughput. We introduce a novel solution to decode temporally interleaved analog signals at nanosecond timescales to achieve high-speed time-multiplexed imaging. Detailed characterization and comparison of the modified and the original two-photon microscopy system was performed.

**OCIS Codes:** (190.4180) Multiphoton processes; (180.5810) Scanning microscopy; (170.2520) Fluorescence microscopy.

## 1. Introduction

Two-photon laser scanning microscopy (TPLSM) has become the standard non-invasive tool for *in vivo* functional recording of calcium signals linked to neural activity [1]. Conventional microscopes equipped with specialized two-photon microscopy objectives typically allow imaging within a field-of-view (FOV) of 1 mm. The emergence of advanced mesoscopic imaging systems with custom-designed optics and a FOV of 3 – 5 mm [2, 3] provides access to nearly 100× larger brain volume for functional imaging, and has opened new frontiers for studying *in vivo* brain function and information processing across multiple cortical areas in small animals. Due to limited imaging speed, researchers must restrict the size of an imaging area, the number of regions of interest (ROIs), and the number of axial planes, such that each plane is imaged with satisfactory temporal resolution. Faster calcium imaging techniques are highly desired to further expand the scope of opto-physiologal studies of brain function. To address this need, a variety of methods have been introduced in recent years to achieve higher imaging throughput [4]. Select examples include imaging with Bessel beams [5, 6], engineered point spread function (PSF) (aka sculpted light) [7], targeted path galvanometer scanning [8], dual-axis two-photon imaging [9], and light sheet illumination [10]. Multiplexed two-photon imaging methods enabling simultaneous multi-site recordings with multiple excitation beams and a single detector are emerging as well [3, 11–14]. These approaches include time-division multiplexing (TDM) [3, 11, 12], frequency-division multiplexing (FDM) [15–17], binary encoding [18], and numerical image analysis [14]. These methods, however, have not achieved a widespread usage and remain implemented by only a few select research groups, possibly due to their technical complexity. To gain imaging speed, each of these methods inevitably accepts different forms of tradeoffs in either one or more key parameters, including imaging resolution, signal-to-noise ratio (SNR), excitation power, and imaging depth.

A custom two-photon random-access microscopy (2P-RAM) system recently developed by Flickinger *et al*. [3] and commercialized by Thorlabs as Multiphoton Mesoscope (MM) combines a number of advanced imaging methods and technical solutions to set new performance benchmarks in two-photon imaging. In addition to a large FOV of 5 mm, the systems utilizes principles of remote focusing [19, 20] to achieve nearly aberration-free imaging within an optically accessible volume. The system features rapid 3D positioning of the scanning beam within 10 ms or less, thus enabling flexible selection of specific ROIs in a volume that can be imaged sequentially for the duration of an experiment. The system features an optimized two-lens condenser similar to the one described previously [21] for efficient photon collection. With its 12 kHz resonance galvanometer scanner, the MM achieves a frame rate of ∼ 44 Hz while imaging an area of 600 µm × 600 µm, and can sequentially image 4 focal planes at a temporal resolution of ∼ 10 Hz to capture calcium transients.

In this work, we introduce a modified MM configuration, the Dual-Plane Multiphoton Mesoscope (MM^2x^), which effectively more than doubles the imaging throughput of the original system. We utilize a previously unused excitation pathway orthogonally polarized with respect to the existing one to add a second imaging channel. This additional excitation path uses its own remote focusing unit responsible for axial positioning of the imaging plane, so that two planes can be positioned independently in axial direction. Both channels use the same opto-mechanics for lateral positioning and scanning. We utilize a TDM approach to encode excitation laser pulse trains with a temporal delay and decode temporally interleaved fluorescence signals from each channel based on their arrival time at the detector. As a result, we achieved simultaneous imaging from two focal planes independently positioned in axial direction. Furthermore, we performed a detailed characterization of the original MM and the modified MM^2x^ to evaluate and compare their performance. While we describe modification of a specific TPLSM system, the MM, the same principles and technical solutions can be easily implemented to enable multiplexed imaging and enhanced performance of any conventional TPLSM system.

## 2. Methods

Mechanical design was performed with SolidWorks (Dassault Systèmes). Time-resolved electric signals were recorded with a fast oscilloscope (204MXi-A, LeCroy) at sampling rates of 5 GS/s or 10 GS/s. Custom MATLAB (MathWorks) routines were used to analyze the recorded data. Neuronal activity was recorded in a Slc17a7-IRES2-Cre;Camk2a-tTA;Ai93 mouse, in compliance with the Allen Institute’s animal imaging protocols. Motion-correction was performed with the ImageJ plugin moco [22]. In test experiments we imaged fixed brain slices labeled with GCaMP6f and pollen grain slides (Carolina Biological) stained with Fast Green FCF.

## 3. System design

A diagram of the imaging system is shown in Fig. 1a. The original MM system was modified to include the following components: a second electro-optical modulator (EOM), an additional pathway for the orthogonally polarized beam with a temporal delay of 6.25 ns, a second remote focusing unit identical to the original one, and demultiplexing electronics. Two EOMs positioned in series achieve complete excitation power control in both imaging planes and allow for the most efficient use of available laser power. The first EOM controls the total excitation power. The second EOM controls the power splitting ratio between both beams by rotating the polarization of the incoming laser beam before passing through a polarizing beam splitter (PBS). This treatment of the incoming beam, consisting of a series of ultrashort pulses, produces two orthogonally polarized beams. Delaying one relative to the other by 6.25 ns and recombining these beams with another PBS effectively creates two interleaved temporally encoded excitation pulse trains. These beams are directed to the input PBS of the dual-plane remote focusing unit which first separates the orthogonally polarized beams. The separated beams are steered into two independent remote focusing assemblies, where beam divergences are controlled to focus these beams in different focal planes, then recombined again and directed to the XY scanners. In this configuration, focal planes are positioned independently in axial dimension while remaining coupled laterally. PMT signals (H11706–40, Hamamatsu) were amplified with a trans-impedance amplifier (HCA-400M-5K-C, Femto, bandwidth 400 MHz) and directed to a custom demultiplexing circuit. A PMT gain setting of 0.65 V was used in all experiments. Temporal alignment of optical signals from PMT and control signals was performed with an adjustable coaxial delay line (DB64, Stanford Research).

**Fig. 1.**
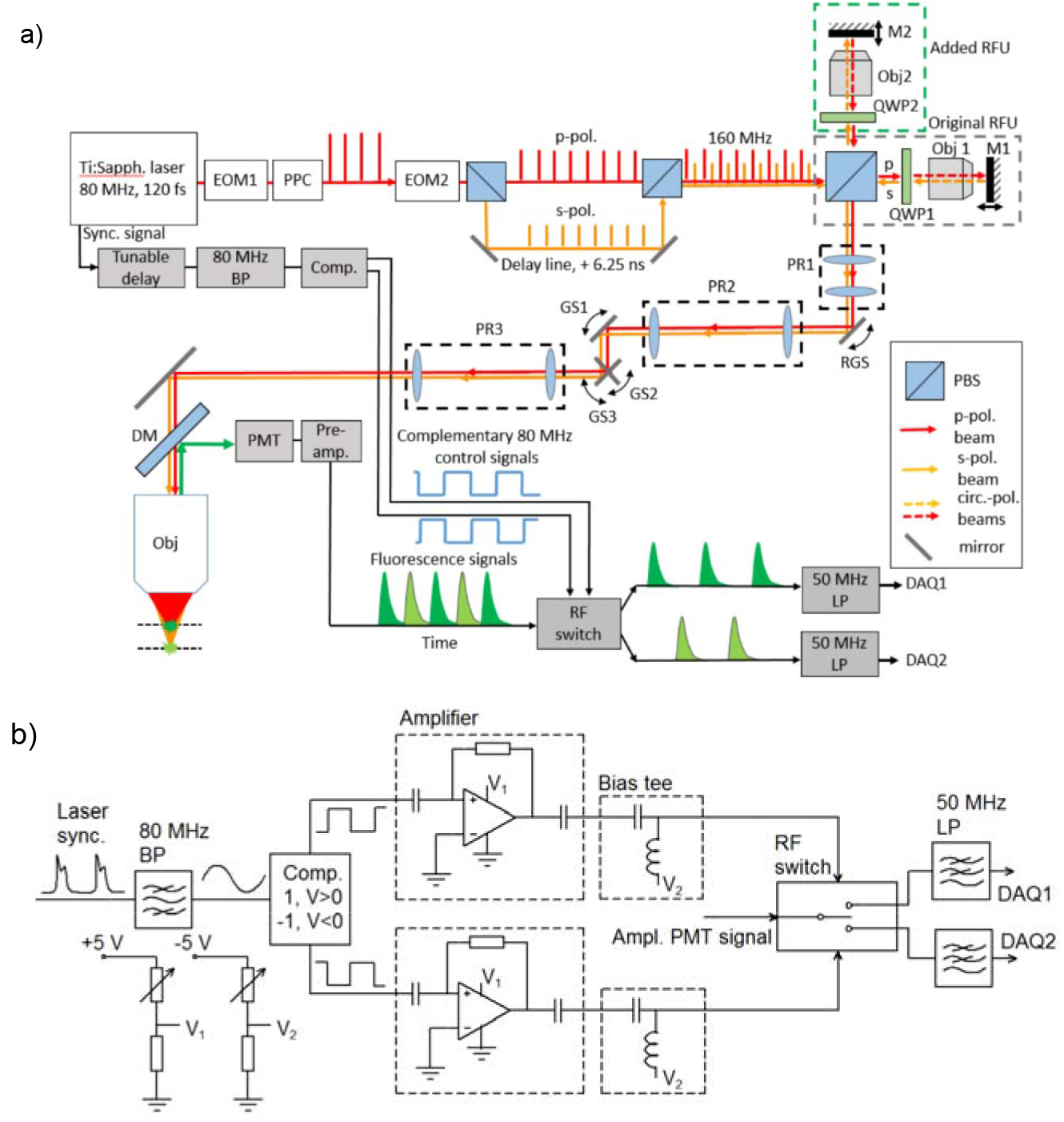
(a) System diagram of Dual-Plane Multiphoton Mesoscope (MM^2x^) imaging system with temporally multiplexed excitation and emission. Mirrors M1 and M2 control axial positioning of the corresponding focal planes. Abbreviations: EOM – electro-optical modulator, PPC – pulse prism compressor, Obj – objective, RFU – remote focusing unit, M – mirror, QWP – quarter wave plate, PR – pupil relay, RGS – resonance galvanometer scanner, GS – galvanometer scanner, DM – dichroic mirror, PMT – photomultiplier tube, BP – band-pass filter, LP – low-pass filter, DAQ – data acquisition channel. (b) Electrical diagram of the signal demultiplexing circuit.

A schematic of the signal demultiplexing circuit is shown in Fig. 1b. Our design is based on a radio-frequency (RF) switch (CMD196C3, Custom MMIC). Temporal demultiplexing requires synchronization with the laser pulse frequency which is not stationary and dithers in time. We utilized an 80 ± 2 MHz bandpass filter (BPF) (3016, KR Electronics, Inc.) and a comparator (LTC6957-HMS3, Analog Devices) to derive complimentary square wave signals with 50% duty cycle from the laser synchronization signal. The comparator output signals are scaled to a peak-to-peak amplitude of 5 V with a high-bandwidth amplifier (GVA-83+, MiniCircuits) and shifted from the common mode V_cm_ = 0 V to V_cm_ = –1 V using a bias tee (ZFBT-4R2GW+, MiniCircuits). Note, the switch controls require negative voltages, and excessive positive voltages may damage the integrated circuit. The control inputs of the RF switch were terminated with 100 Ohm resistors. After demultiplexing, the two high-bandwidth signals were passed through 50 MHz low-pass filters (BLP-50+, Mini-Circuits), directed to the digitizer inputs (NI FlexRIO, National Instruments) and sampled at 80 MHz. The original MM control software (ScanImage, Vidrio LLC) was customized by Vidrio to provide controls for both RFUs and EOMs and accommodate random-access imaging at two focal planes.

Fig. 2 shows CAD models of original MM and modified MM^2x^ system with the dual-plane remote focusing assembly installed. The dual-RFU module was designed as an attachment which could be easily added to and removed from the system without modifying any existing mechanical components. We designed an intermediate mounting plate providing anchor points for all additional hardware, while the plate was attached to the MM breadboard using existing fixation points. Additional components, including mounting brackets, redesigned tip-tilt stages and voice-coils of both RFUs were fixed on this plate (see Fig. 2b). Because of limited space within the MM enclosure, the s-polarized, second excitation beam was reflected orthogonally to the breadboard towards the second remote focusing unit using a 45° prism mirror. This way, the dual-plane RFU assembly fits inside the MM enclosure without restricting the system’s XYZ and tilt adjustments.

**Fig. 2.**
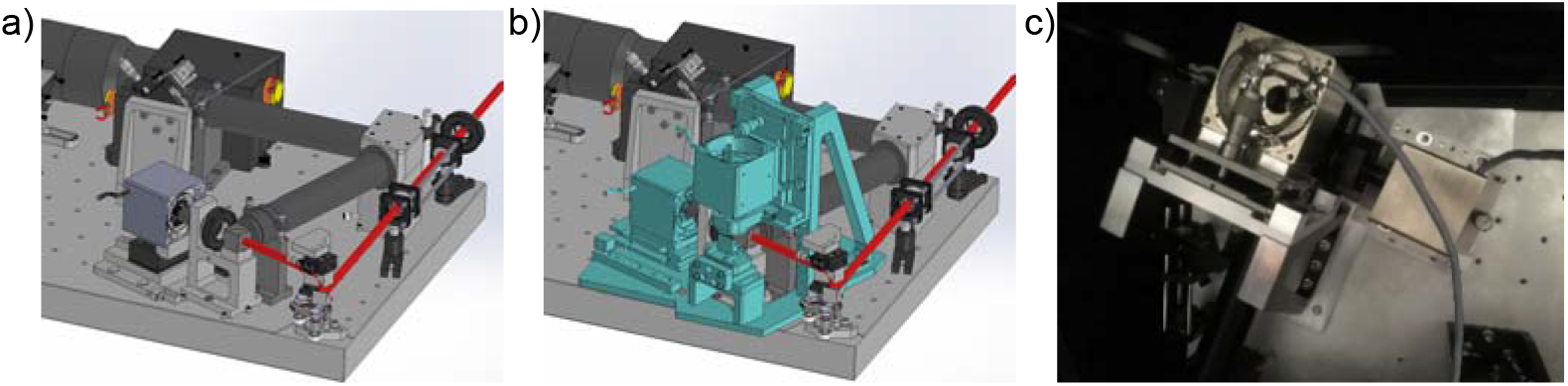
(a,b) CAD models of original MM (a) and modified MM^2x^ (b). Added components are highlighted in cyan color. (c) Top view of installed dual-plane remote focusing assembly.

## 4. System characterization

We evaluated and compared the optical performance of our modified MM^2x^ to the original MM by measuring cross-talk levels between imaging channels, signal dynamic range, noise levels of acquired images, and PSFs at different FOV regions and depths. The modular design of the MM^2x^ allowed us to easily switch between configurations by simply removing the demultiplexing electronics from the signal detection path, leaving current amplifier and low-pass filter in place.

Time-interleaved excitation pulses with a combined pulse rate of 160 MHz generate a corresponding sequence of fluorescence signals arising from different locations. This pulse rate defines the width of the signal integration window, i.e., 6.25 ns. Therefore, the demultiplexing circuit must have sufficient bandwidth to separate signals from both channels. Fig. 3a illustrates the operation of our custom demultiplexing electronics. An input square wave of 0 to 1 V and “on” duration of a 100 ns is toggled between two DAQ channels by control signals derived from the laser sync signal. Both output signals have a period of 12.5 ns and a duty cycle of 39%., i.e. “on” time relative to a signal period. The output signal amplitude gets attenuated by 1.6 ± 0.2 dB, consistent with the specified insertion loss. Fig. 3b shows the same signals on an expanded time scale. The observed 10–90% rise/fall time of these signals was as low as 0.6 ns. While high bandwidth is important for the TDM method, it is also essential to consider the duration of the laser pulse-induced fluorescence signals. The fluorescence lifetime τ of calcium indicators based on green fluorescence protein appears in the range of 2.7 – 3.2 ns [23, 24]. This lifetime is sufficiently long to extend the fluorescence signal into the next integration window, thereby producing significant cross-talk between imaging channels. Averaged PMT signals corresponding to a single photon detection event and measured time-resolved fluorescence signals of GCaMP6f and pollen grains (PGs) are shown in Fig. 3c,d. The impulse response traces show the main peak with a FWHM of ∼2.5 ns as defined by the preamplifier bandwidth, and a smaller side lobe, which is due to the reflection of broadband pulses at the PMT-to-preamplifier cable (length ∼ 5 cm). The fluorescence signal of PG in Fig. 3d is noticeably shorter. The secondary peak is the reflected signal, which appears further away from the main peak due to a longer lead cable (length ∼ 45 cm). Note, time-resolved measurements with PGs were performed before reducing the lead cable length, while the other experimental data presented here were not affected by the delayed RF reflections. Clearly, fluorescence signals from a single channel exceed in time the theoretical 6.25 ns interval, thus, the late portion of them appear in the next integration window causing channel cross-talk. Here, we define cross-talk as the fraction of a ghost signal with respect to the signal of interest, when excitation present in only one channel.

**Fig. 3.**
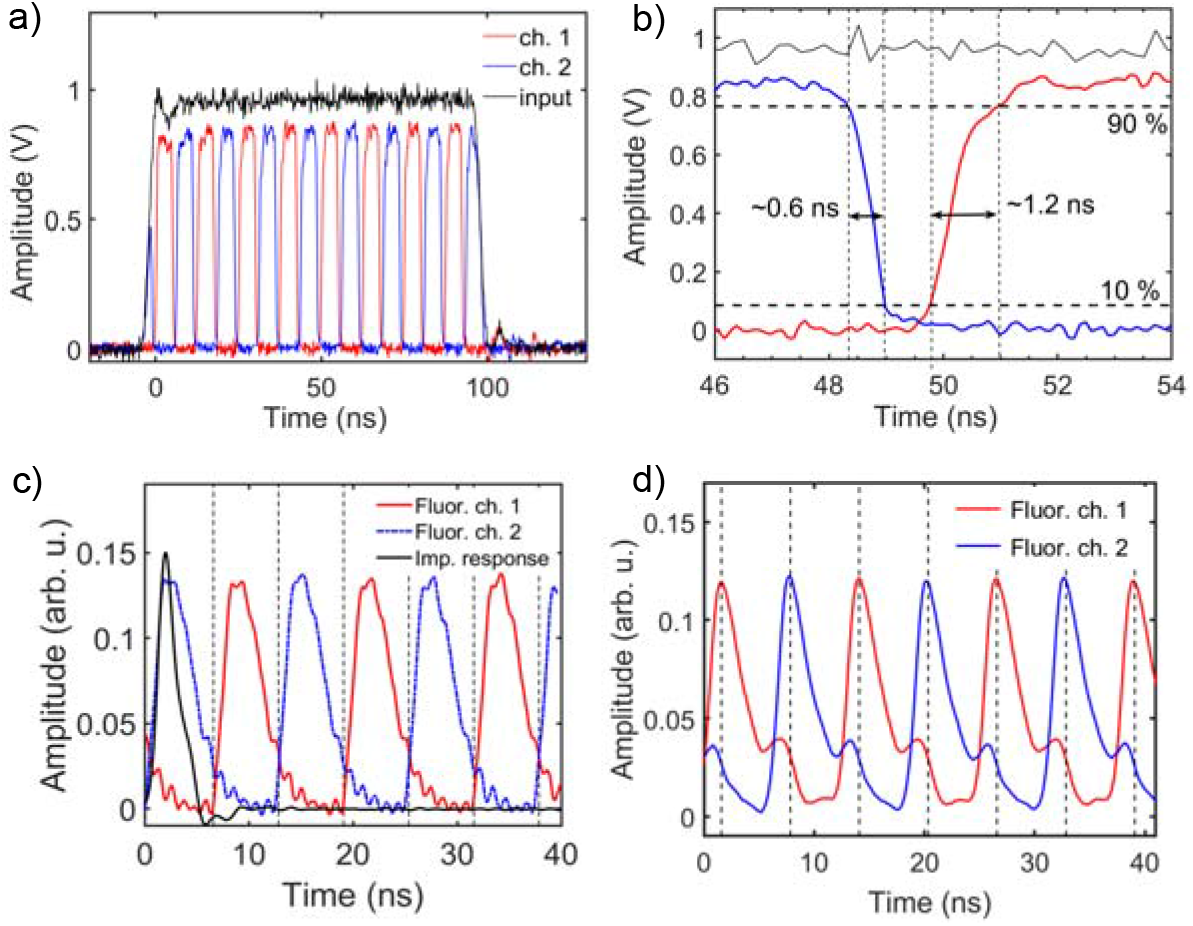
Operation of demultiplexing electronics. (a) Input and output signals recorded at demultiplexing circuit. (b) Time-expanded section from (a). (c) Time-resolved impulse response of amplified PMT signal, averaged over individual photon detection events, and time-resolved GCaMP6f fluorescence profile. (d) Time-resolved fluorescence profile of pollen grains, interval between vertical grid lines 6.25 ns, signals are averaged 8,000 times.

Fig. 4a,b shows pairs of motion-corrected and time-averaged images of mouse brain in two channels recorded simultaneously at depths of ∼200 – 300 °m, while exciting the samples only in one of the channels. Thus, each set of images shows fluorescence signals from cells in the plane of interest, and ghost images of the same cells due to the presence of cross-talk. Note however, pixel intensities in the ghost images in Fig. 4 are scaled up by a factor of 10 for visibility. We extracted calcium signals from selected cells to demonstrate their relative intensities in both imaging channels (see Fig. 4c,d). Fig. 4e shows cross-talk values computed as ratio of the measured calcium signals. The average cross-talk values computed from traces in Fig. 4e are 5.6 ± 0.7% for Ch. 1 → Ch. 2 and 7.0 ± 1.1% for Ch. 2 → Ch. 1. Due to the stochastic nature of the fluorescence and the measurement noise, crosstalk values are expected to fluctuate from frame to frame.

**Fig. 4.**
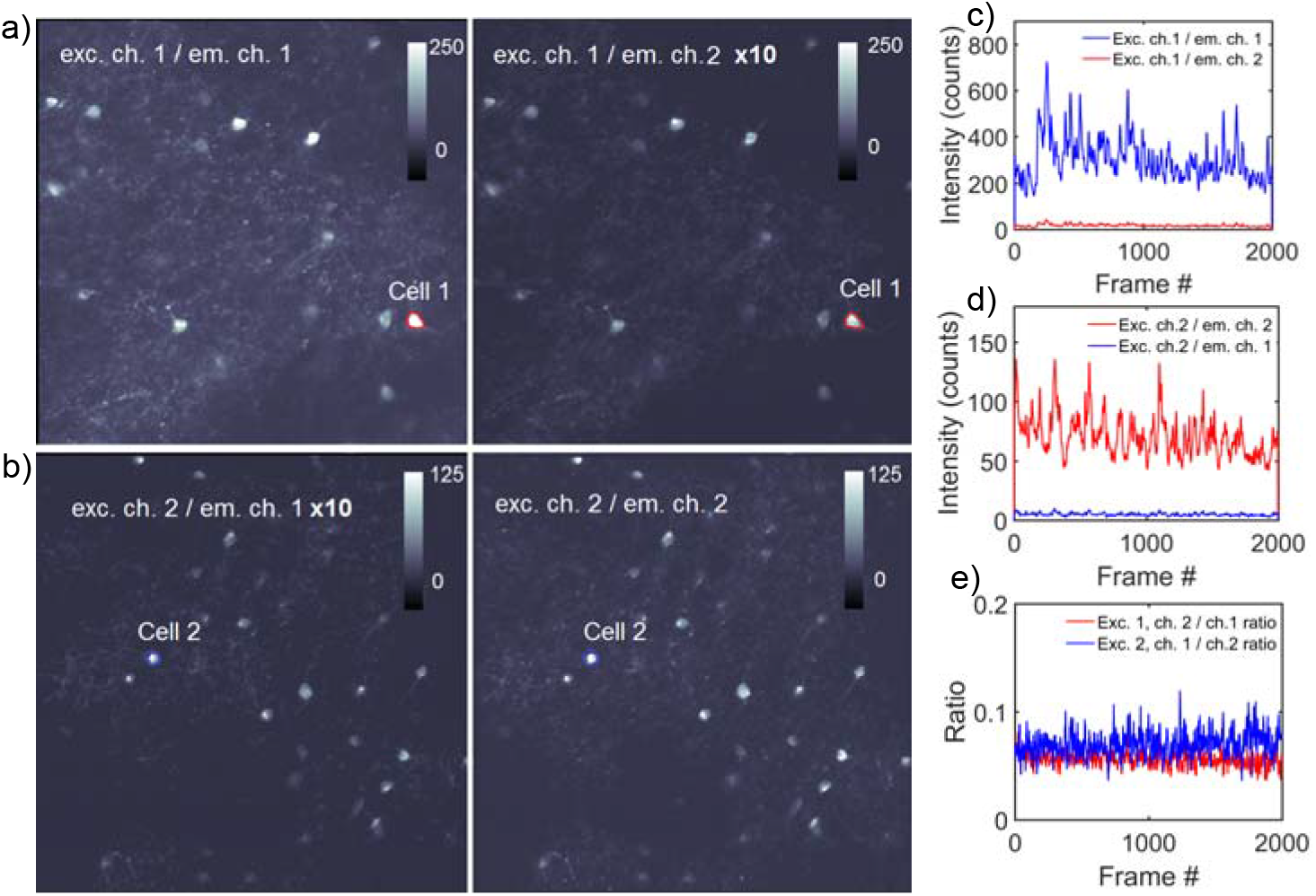
(a,b) Channel cross-talk evaluation. *In vivo* images of mouse cortex in two simultaneously recorded imaging channels using excitation in either first or second imaging channel only. Images were averaged 2,000 times. Note the scaling factor of non-excited channels. (c,d) Calcium signals extracted from cell bodies shown in (a,b), respectively. (e) Channel cross-talk of calcium signals from (c,d).

We compared the performance of both original MM and modified MM^2x^ systems by recording images of test samples under the same experimental conditions. The axial positions of both channels were adjusted to acquire images from the same focal plane. Examples demonstrating qualitative equivalence of these systems are shown in Fig. 5a. Images of the same region were recorded at different excitation power levels. The corresponding average fluorescence signals from a single PG indicated by the arrow in Fig. 5a are shown in Fig. 5b. Below 4,000 counts, signal levels in both channels are nearly the same in both MM and MM^2x^. Mean-variance plots computed from all recorded images from MM and MM^2x^ demonstrate almost the same SNR in both imaging systems below fluorescence saturation threshold (see Fig. 5c).

**Fig. 5.**
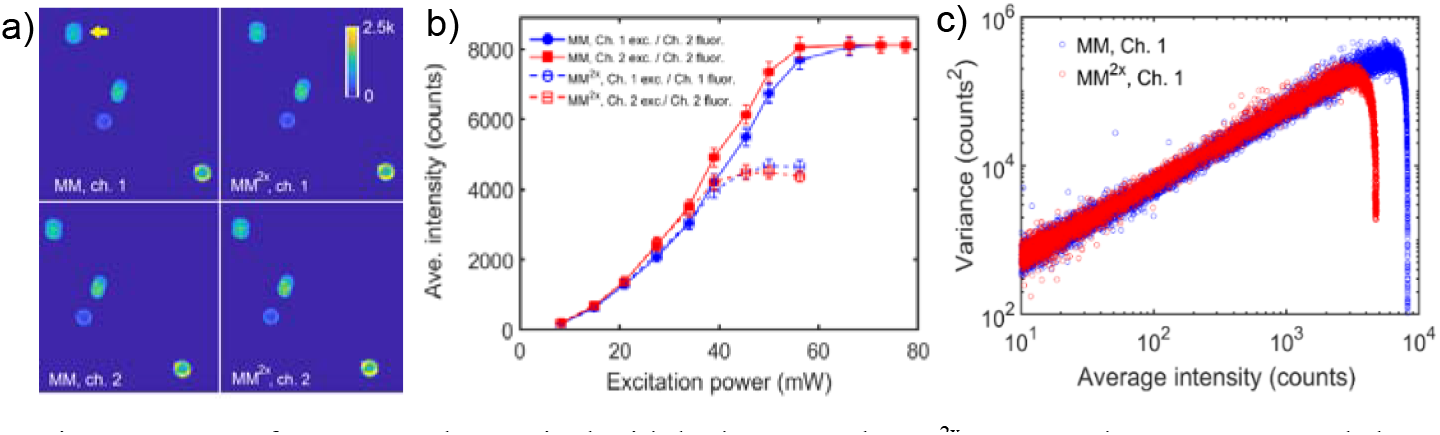
Images of a PG sample acquired with both MM and MM^2x^. Images shown were recorded at an excitation power of 28 mW and averaged 100 times. (b) Average fluorescence signal intensity of the pollen grain indicated by the arrow in (a) at different excitation power levels. (c) Mean-variance plot computed from all recorded PG images in channel 1 of both MM and MM^2x^ systems. For clarity only data from channel 1 are shown.

A similar set of experiments was conducted using a fixed brain slice, the results are presented in Fig. 6. Fluorescence signals appear slightly attenuated in the images recorded with the modified system as compared to the original one, with attenuation coefficient of 0.76 in channel 1, and 0.88 in channel 2. We note that biological samples are significantly less photostable, and photobleaching effects could have affected the measurement of fluorescence signal in MM, ch. 2 (filled red circles), since it was measured last. Mean-variance plots computed from these images are shown in Fig. 6d. Again, we observed nearly the same SNR values in both imaging systems.

**Fig. 6.**
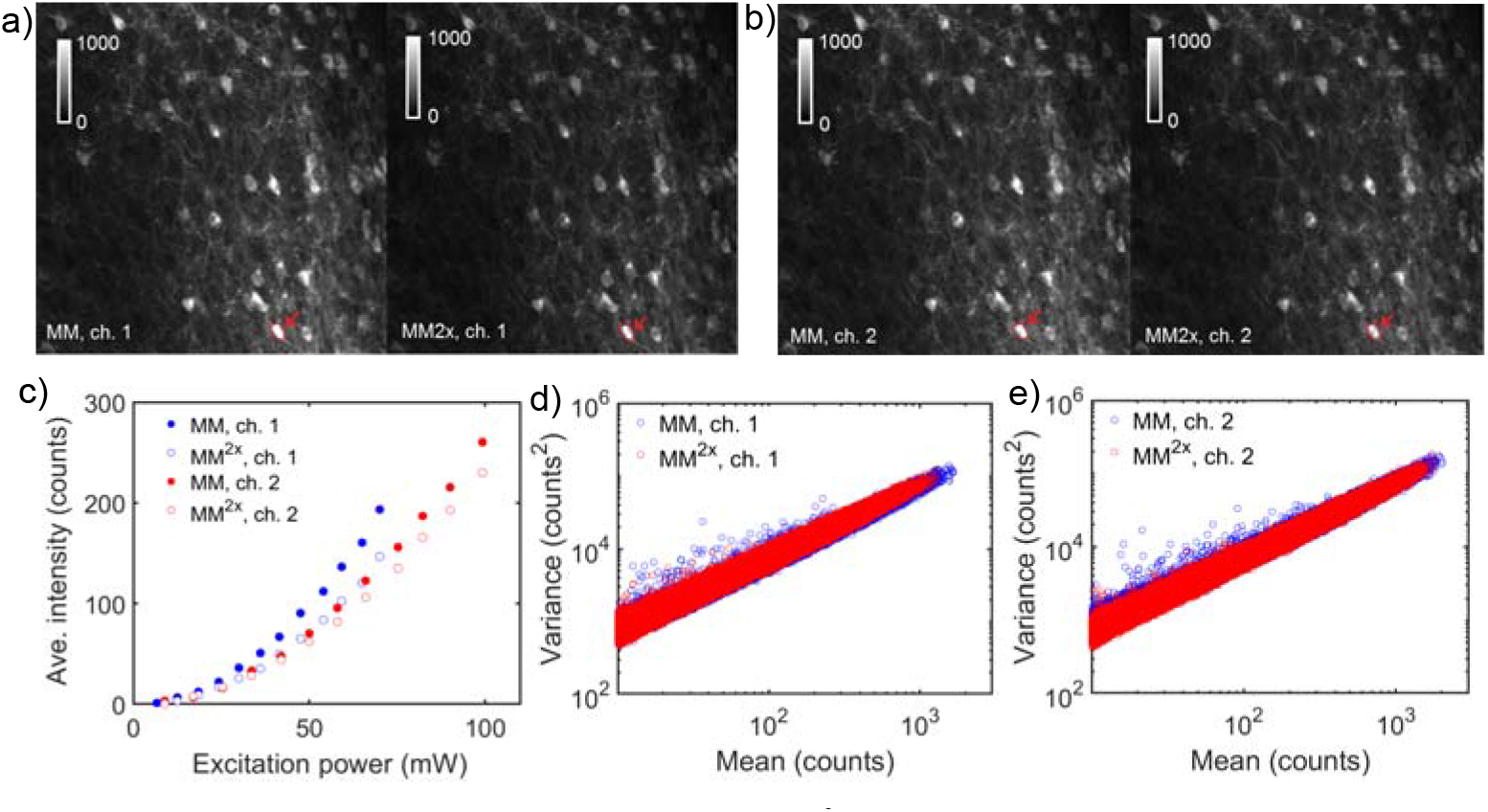
Fixed brain slice imaged with both MM and MM^2x^, averaged 400 times. The images from Ch. 1 and Ch. 2 were recorded at 70 mW and 82 mW excitation intensity, respectively. (c) Average fluorescence signal of the cell body indicated by the arrows in Figs. 6a,b at different excitation intensities. (d,e) Mean-variance plots computed from corresponding unaveraged image series.

Fluorescence signals of neurons and PGs were recorded while adjusting the electronic delay line, thereby changing the relative position of control signals with respect to fluorescence signals. As before, excitation was present in only one channel. Fig. 7 shows the normalized fluorescence intensity profiles as a function of time delay. As expected, we observed a periodically varying signal in both imaging channels as a function of temporal offset. In Fig. 7a, the optimal delay providing the highest contrast was 4.5 ns, with a crosstalk of ∼ 7%. Cross-talk level in PG sample (Fig. 7b) was negligible, less than 1 %.

**Fig. 7.**
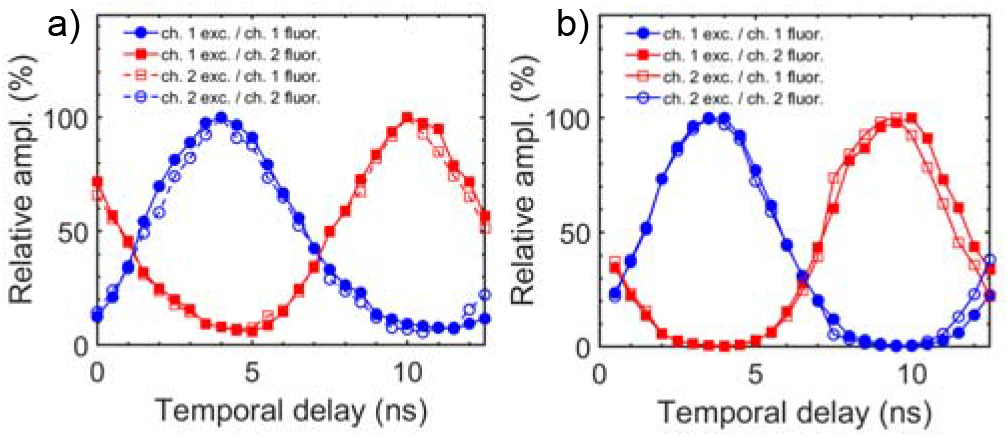
Comparison of normalized average emission intensities in two imaging channels as function of time delay. Profiles in (a) and (b) correspond to fluorescence signals of GCaMP6f-labeled cells and PGs, respectively.

We measured PSFs across the imaging volume by acquiring z-stacks of 200 nm fluorescent beads immobilized in a 1% aqueous agarose gel matrix at different imaging depths from 0 μm to 500 μm, and different regions in the FOV. Examples of recorded PSFs are shown in Fig. 8. Overall we observed nearly identical PSFs in both imaging channels, similar to the previously reported values [2].

**Fig. 8.**
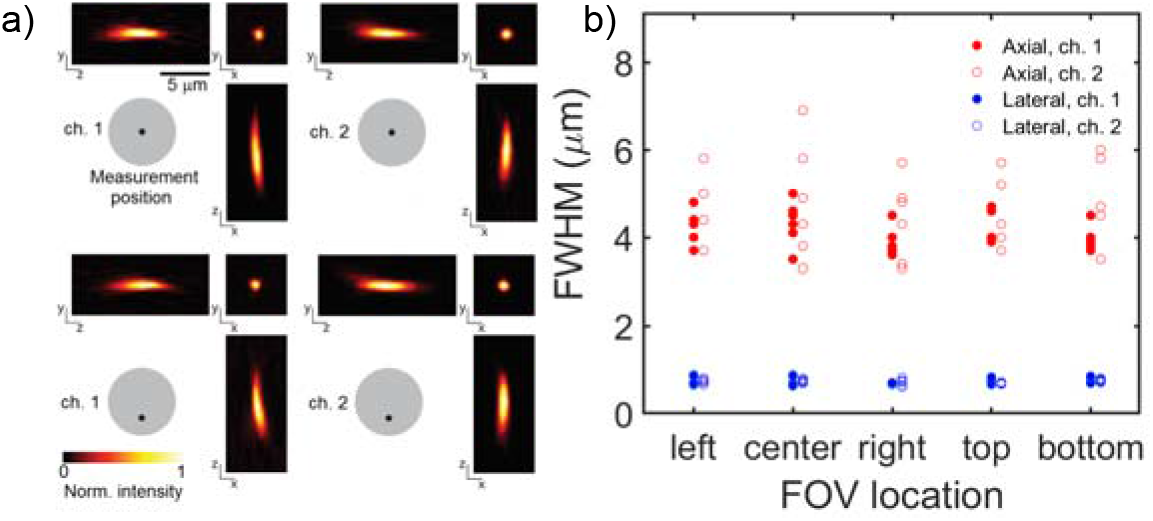
(a) Maximum intensity projections of PSF. The PSFs shown were measured at a depth of 300 μm. (b) PSF full width at half maxima (FWHM) in both imaging channels, recorded at depths of 0, 100, 200, 300, 400, 500 μm, and different locations within the FOV.

## 5. Discussion

Although application of TDM in two-photon imaging was demonstrated years ago [11], this method has not yet evolved to the level of commercial imaging instrumentation. Up to date, only a few research groups in the two-photon imaging community have implemented this technology [3, 11, 12]. Demultiplexing of time-interleaved high-bandwidth PMT signals at rates of ca. 100 MHz is technically challenging. There are two critical issues to be resolved. The first task is synchronizing the detection electronics with the non-stationary laser repetition rate which is critical to correctly assign signals to imaging channels. Since laser pulse rates depend on the resonator, any path length change, for example, due to alterations in ambient temperature, will affect this rate. One may recall that a periodic sequence of electronic pulses generated from an optical laser pulse train is represented by a discrete set of frequencies centered at the main frequency. Thus, it is possible to isolate the main frequency component using an appropriate 80 MHz RF filter, and convert the resulting sine wave into a square waveform, and two complementary square waveforms using an RF comparator circuit (see Fig. 1b). The second and most challenging task is signal demultiplexing itself. Electrical signals at the PMT output corresponding to neural activity are largely diverse, ranging from isolated high-bandwidth spikes arising from individual photon detection events to significantly higher and longer bursts from high-count photon fluxes. Overall, the dynamic range of digitized signals spans more than 4 orders of magnitude. The resulting requirements for the demultiplexing electronics in terms of detection sensitivity, detection bandwidth, and dynamic range are difficult to meet. Published reports each describe their own demultiplexing scheme [3, 11, 12]. Unfortunately, a detailed system characterization and comparison with a conventional TPLSM setup measured in identical experimental conditions were usually not provided, and it remains unclear to what extent the performance of the published imaging systems was degraded by using TDM. For example, our initial demultiplexing circuit was based on an analog multiplier ADL5391 (Analog Devices) as in [11]. While this design achieved demultiplexing (data not shown), it resulted in a more than 5-fold increase in background noise due to the presence of additional amplification stages and consequently significant background shifts in case of large intensity high-bandwidth signals. Therefore, we revised our design, and our current demultiplexing circuit is based on a fast RF switch with a manufacturer-specified switching time of 1.8 ns. While it is recommended to apply control voltages from 0 to –5 V, we even achieved faster switching dynamics by slightly shifting the complementary 80 MHz square wave controlling the switch operation towards positive voltages. We observed a lag time of ∼ 1 ns, followed by ∼ 0.6 – 1.2 ns rise/fall time after stepping the control signals. This lag effectively resulted in a duty cycle reduction of 39% of the signal integration temporal interval (see Fig. 3a).

An important parameter of TDM systems is the loss of useful signal during demultiplexing. Both the fluorescence lifetime of a fluorophore and bandwidth of the detection electronics affect the duration of fluorescence signals induced by single laser pulses. If these signals last longer than a measurement interval, then a portion of them will leak into the next temporal window. Therefore, TDM systems favor molecules with shorter florescence lifetimes, resulting in reduced cross-talk and less losses of useful signal. A direct comparison of fluorescence signals from PG samples imaged with both MM and MM^2x^ is shown in Fig. 5, demonstrating a nearly identical performance of both systems, as long as no signal saturation takes place. A similar comparison of fluorescence signals from GCaMP6f-labeled samples (see Fig. 6) shows a reduction of signal amplitude of up to 24% in the modified system. Such signal loss is expected since the average GCaMP6f fluorescence signal exceeds the 4.9 ns temporal sampling interval, as shown in Fig. 3. Part of it can be attributed to the insertion loss of the demultiplexing electronics, which only changes the amplification factor (or conversion gain) of the system and does not affect the number of detected photons. Using the data from Fig. 3, we estimated the fraction of GCaMP6f fluorescence appearing outside of the integration window and thus lost to be ∼ 17%, while ∼ 5% will end up in the next window. These estimates agree well with experimentally observed cross-talk values in Figs. 4, 7. These losses appear rather small, given that mean-variance plots from both systems in Fig. 7 and 8 appear closely matched, before signal saturation effects takes place. For example, the linear slopes in Fig. 6 from GCaMP6f-labeled images recorded with MM and MM^2x^ in both imaging channels equal 70 ± 2 counts, which is within ± 3% of their average value. The linear slopes corresponding to the data from Fig. 5 (PG samples) are 59 ± 2. The relative difference of these coefficients indicates signal loss in GCaMP6f-labeled samples as compared to samples stained with Fast Green FCF due to the differences in their fluorescence lifetimes.

The signal saturation in MM, which was ∼ 8,000 counts (∼ 2^13^) in Fig. 5b, is caused by the 14-bit DAQ electronics which has a limited input voltage range of ±1 V. Note, the signal reference level is currently set at 0 V. Therefore, signals appearing at the DAQ input must have amplitudes < 1V to avoid non-linearity in the recorded signals. Peak voltages in amplified PMT signals at the given PMT gain setting routinely exceed 1 V, and the use of low-pass filters is essential for matching the signals to both digitization window and rate of the DAQ. With the demultiplexing circuit in place, the signal saturation occurs at approximately 4,000 counts, which corresponds to a 50% reduction in the detection dynamic range. In test experiments, we observed the attenuation factor in the demultiplexing circuit progressively increasing with the amplitude of input signals. The RF switch limits their amplitude to a range of approximately ±2 V. These voltage levels seem to be above the manufacturer-specified operating limits of the high-bandwidth preamplifier. Since the RF switch operation is identical at positive and negative signal polarity, it is possible to extend the dynamic range by introducing a DC offset to the preamplifier output and measuring the signals with respect to a new reference.

The modified MM^2x^ system provides a significant enhancement in the imaging throughput in specific experimental conditions that require sequential imaging of the same area at different depths. For example, considering the simplest experimental configuration of imaging only two stacked planes, the original MM system can achieve an overall frame rate of ∼ 15 Hz, with 22.7 ms frame measurement time and 10 ms axial repositioning time. In the same configuration, MM^2x^ will maintain a frame rate of 44 Hz. The demonstrations of *in vivo* functional imaging at increased frame rates for studying brain function is underway.

## 6. Conclusions

We developed a modification of the Multiphoton Mesoscope which enables simultaneous imaging of two focal planes independently positioned along axial direction. We utilized a previously unused imaging channel to introduce a second excitation beam into the system, orthogonally polarized relative to the original beam, and a second remote focusing unit identical to the original one. Both remote focusing units provide independent positioning of image focal planes, as well as aberration correction of the beam point spread functions. Both beams share the optomechanics controlling lateral beam positioning and scanning. Our upgrade was designed as an add-on to the original system that does not require any modification of the original system components, and which can be installed or removed at will. To enable simultaneous image acquisition, we utilized temporal division-multiplexing, where different excitation beams and corresponding excitation pulses are encoded with a specific temporal delay, and corresponding fluorescence signals detected with a PMT are assigned to different imaging channels based on their arrival time at the detector. For signal demultiplexing, we developed a novel analog solution based on a fast RF switch, operated by control signals derived from the laser synchronization signal. Furthermore, we performed a complete system characterization and compared its performance to the original system. The modified MM^2x^ system achieves an increased imaging throughput of more than a factor of two with only minor degradation in the magnitude of the useful signals.

## Acknowledgments

We wish to thank the founder of the Allen Institute for Brain Science, Paul G. Allen, for his vision, encouragement and support.

